# Benchmarking Large Language Models for Pathogen–Disease Classification in Post-Acute Infection Syndromes

**DOI:** 10.1101/2025.06.30.662395

**Authors:** Syed Mohammed Khalid, Tom Wölker, Leidy-Alejandra G. Molano, Simon Graf, Andreas Keller

**Affiliations:** Clinical Bioinformatics, Saarland University, 66123, Saarbrücken, Germany; Helmholtz Institute for Pharmaceutical Research Saarland (HIPS)–Helmholtz Centre for Infection Research (HZI), Saarland University Campus, 66123, Saarbrücken, Germany

## Abstract

Post-Acute Infection Syndromes (PAIS) are medical conditions that persist following acute infections from pathogens such as SARS-CoV-2, Epstein-Barr virus, and Influenza virus. Despite growing global awareness of PAIS and the exponential increase in biomedical literature, only a small fraction of this literature pertains specifically to PAIS, making the identification of pathogen–disease associations within such a vast, heterogeneous, and unstructured corpus a significant challenge for researchers. This study evaluated the effectiveness of large language models (LLMs) in extracting these associations through a binary classification task using a curated dataset of 1,000 manually labeled PubMed abstracts. We benchmarked a wide range of open-source LLMs of varying sizes (4B–70B parameters), including generalist, reasoning, and biomedical-specific models. We also investigated the extent to which prompting strategies such as zero-shot, few-shot, and Chain of Thought (CoT) methods can improve classification performance. Our results indicate that model performance varied by size, architecture, and prompting strategy. Zero-shot prompting produced the most reliable results: Mistral-Small-Instruct-2409 and Nemotron-70B achieved balanced accuracy scores of 0.81 and 0.80, respectively, along with macro-F1 scores of up to 0.80, while maintaining minimal invalid outputs. While few-shot and CoT prompting often degraded performance in generalist models, reasoning models such as DeepSeek-R1-Distill-Llama-70B and QwQ-32B demonstrated improved accuracy and consistency when provided with additional context.

## 1 Introduction

Traditionally, acute infectious diseases were perceived as short-lived episodes that ended either in patient recovery or death [1]. However, now they are growingly being recognized for their potential to lead to chronic or prolonged symptoms being known as post-acute infection syndromes (PAIS), which refer to the diverse set of persistent conditions following acute infection[1]. Following the COVID-19 pandemic, approximately 65 million individuals experienced ongoing symptoms lasting at least three months post infection [2][3]. Chronic sequelae have been documented following infections with a wide spectrum of pathogens, including SARS-CoV-2, Epstein-Barr virus, Influenza virus, and Ebola virus [4–7]. The nonspecific nature of PAIS symptoms, along with the absence of reliable biomarkers, poses substantial challenges for clinical diagnosis, treatment, and the development of preventive strategies. Although biomedical research output has grown substantially in recent years, only a limited portion of studies have systematically addressed PAIS.[8] As a result, researchers face considerable challenges in identifying clear pathogen–disease associations from the existing largely unstructured literature base. To accelerate discovery in this field, there is a pressing need for scalable, systematic approaches that can synthesize existing knowledge and uncover novel insights beyond the limitations of manual and labor-intensive methods.

Early attempts to automate this task relied on sentence-level co-occurrence. An ontology-based text-mining study extracted 3,420 pathogen–disease pairs from 1.8 million open-access articles [9], reporting 64% precision, 89% recall, and an F-score of 0.74. However, the system accepted only associations occurring at least ten times and with a Normalized Point-wise Mutual Information (NPMI) ≥0.2, thereby discarding many rare but clinically relevant links. Additionally, sentence-level co-occurrence further introduced false positives, often due to abbreviation ambiguities [9]. Such rigid thresholds are especially problematic in contexts where relevant pathogen–disease links are often reported using heterogeneous and evolving terminology.

Generative artificial intelligence (AI) systems have raised considerable interest due to their rapidly expanding capabilities and significant potential across a diverse range of applications[10]. Among these, large language models (LLMs) particularly foundation models built on transformer architecture[11] and reinforcement learning [12], such as GPT-4o from openai[13], Llama 3[14] from Meta, or reasoning models like DeepSeek-R1[15], have emerged as particularly influential, demonstrating impressive performance in diverse tasks such as comprehension [16], summarization [17], text-classification [18], and information retrieval[19].

Despite their observed general capabilities, questions remain regarding the effectiveness of LLMs within the biomedical research domain. Past research presents a conflicting set of approaches and outcomes. On one hand, domain-specific models such as BioBERT [20], when fine-tuned on biomedical datasets, have consistently outperformed generalist LLMs like GPT-2 and Flan-T5 in tasks such as biomedical text classification and reasoning [21, 22]. On the other hand, some studies suggest that generalist models, when coupled with innovative prompting techniques, can rival or even surpass the performance of specialized models [23, 24].

Prompting, in the context of language models, refers to the input crafted to steer the model’s generated output. A model’s performance on a given task can be significantly affected by the way the prompt is formulated, sometimes in unexpected or surprising ways [25]. A wide range of prompting approaches have been explored, demonstrating significant potential in improving LLMs’ performance in biomedical-related tasks [26]. For example, advanced prompting methods such as few-shot learning and retrieval-augmented reasoning have been proven effective in biomedical literature analysis, particularly for entity recognition tasks [21, 27, 28]. In contrast, in structured biomedical classification tasks, simple prompting strategies have often out-performed more complex reasoning techniques such as chain-of-thought (CoT) or retrieval-augmented generation (RAG)[29]. Similar observations were made in other studies, where incorporating external evidence into prompts was found to negatively impact model accuracy [30].

Despite extensive research in biomedical language processing, most existing studies focus on general biomedical tasks or conventional disease contexts. Notably, to our existing knowledge, there is a lack of benchmarking efforts evaluating LLMs for identifying pathogen–disease relationships, particularly within the unique literature of PAIS. To address this gap, we systematically benchmarked state-of-the-art large language models (LLMs) to assess their reliability in accurately identifying pathogen–disease associations in PAIS-related biomedical literature. We employed various prompting strategies, including zero-shot, few-shot, and chain-of-thought (CoT) methods, to evaluate and compare models ranging from small language models (SLMs) to those with 70 billion parameters. Additionally, we analyzed performance differences between generalist foundation, reasoning, and domain-specific models.

## 2 Methods

### 2.1 Dataset

Given the absence of publicly available labeled datasets specifically focused on pathogen–disease relationships within the context of post-acute infection syndromes (PAIS). To address this, we constructed our evaluation dataset tailored to this task. We began by first compiling lists of pathogens, diseases, and their relationships from PathoPhenoDB[31], Disbiome[32], gcPathogen[33], and DO[34]. Using these sources, PubMed queries were created. Any pathogen–disease combination included in a relationship source or that was “similar” was excluded from the queries. We define similarity as either the pathogen or the disease term being contained within the other (e.g., “porcine diarrhea virus” and “diarrhea”), or having an edit distance of 0.85 or greater (e.g., “Leishmania” and “leishmaniasis,” where the larger term was truncated by three characters to set a stricter threshold).

Next, the query results were refined by keeping only the “lowest rank” pathogen for each abstract. For example, if the same abstract returned both “Rickettsia” and “Rickettsia africae,” only “Rickettsia africae” (the more specific term) was kept. From this refined set of results, we selected 1,000 entries randomly and uniformly. Finally, the labels for each entry in the dataset were assigned manually according to the following specified set of rules (An abstract was labeled as indicating a relationship if it fulfilled any one of Criteria 1–4):

Abstract Evaluation Criteria

1. Does the abstract investigate the pathogen and report evidence for the disease?
2. Does the abstract investigate the disease and report evidence for the pathogen?
3. Does the abstract focus on the relation between the Pathogen and the Disease?
4. Does the abstract state the relation between Pathogen and Disease, but does not focus on it?

We used 994 of the samples for evaluation and reserved 6 for few-shot prompting. The dataset consists of 70% negative and 30% positive examples and includes a broad range of pathogen and disease types.

### 2.2 Models

We have selected a wide range of LLMs to evaluate their effectiveness in classifying pathogen-disease relationships within biomedical literature. We limited our study to open-weight checkpoints released that are hosted on Hugging Face and runnable on 2 ×NVIDIA H100 80 GB or less. Our selection covers multiple scales and architectures, structured as follows (Table 1):

**Table 1.** LLM Specifications.

| Model | Parameters | Domain | Description |
| --- | --- | --- | --- |
| Llama-3.1-70B-Instruct | 70B | General | Decoder-only, Instruction tuned generative model. |
| Llama-3.3-70B-Instruct | 70B | General | Decoder-only, Instruction tuned generative model. |
| Llama-3.1-Nemotron-70B-Instruct | 70B | General | Decoder-only, Instruction tuned generative model. Fine-tuned Llama-3.1-70B-Instruct with the Llama-3.1-Nemotron-70B-Reward and HelpSteer2-Preference prompts. |
| Qwen2.5-72b-Instruct | 72B | General | Decoder-only, Instruction tuned generative model. |
| Mistral-Small-Instruct-2409 | 22B | General | Decoder-only, Instruction tuned generative model. |
| Phi-3.5-MoE-Instruct | 42B | General | Mixture-of-expert model, Decoder-only, Instruction tuned generative model. |
| Llama-3.1-8B-Instruct | 8B | General | Decoder-only, Instruction tuned generative model. |
| Meta-Llama-3-8B-Instruct | 8B | General | Decoder-only, Instruction tuned generative SLM. |
| Qwen2.5-7B-Instruct | 7B | General | Decoder-only, Instruction tuned generative SLM. |
| Ministral-8B-Instruct-2410 | 8B | General | Decoder-only, Instruction tuned generative SLM. |
| Phi-4-Mini-Instruct | 4B | General | Decoder-only, Instruction tuned generative SLM. |
| QwQ-32B | 32B | General | Reasoning model from the Qwen series. Compared to conventional instruction-tuned models, QwQ excels at thinking and reasoning. |
| DeepSeek-R1-Distill-Llama-70B | 70B | General | Decoder-only, built on Llama-3.3-70B-Instruct. |
| DeepSeek-R1-Distill-Llama-8B | 8B | General | Decoder-only, built on SLM Llama-3.1-8B. |
| DeepSeek-R1-Distill-Qwen-7B | 7B | General | Decoder-only, built on Qwen2.5-Math-7B. |
| DeepSeek-R1-Distill-Qwen-32B | 32B | General | Decoder-only, built on Qwen2.5-32B. |
| BioMistral-7B | 7B | Bio | Decoder-only, built on Mistral and further pre-trained on PubMed Central. |
| PMC-LLaMA | 13B | Bio | Biomedical LLM continually pre-trained on a large corpus of 4.8M biomedical papers |
| Meditron-70B | 70B | Bio | Decoder-only, Built on Llama-2, pre-trained on a comprehensively curated medical corpus, including selected PubMed articles, abstracts, and internationally-recognized medical guidelines |

#### 2.2.1 Large Generalist Foundation Models

We include models from Meta’s Llama series, namely Llama-3.3-70B-Instruct(hereafter referred to as Llama-3.3-70B) and Llama-3.1-70B-Instruct(Llama-3.1-70B), which are 70-billion-parameter, instruction-tuned generative models known for their strong generalist performance across benchmarks[35]. Additionally, we included NVIDIA’s Llama-3.1-Nemotron-70B-Instruct(Nemotron-70B), a reinforcement learning from human feedback (RLHF) tuned variant of Llama-3.1-70B[36]. Furthermore, we include Qwen2.5-72B-Instruct(Qwen-72B) from the Qwen series, another strong performer across various benchmarks [37].

#### 2.2.2 Mid-sized Generalist Foundation Models

We intend to evaluate models in the intermediate parameter range, specifically Phi-3.5-MoE-Instruct(Phi-3.5-MoE) and Mistral-Small-Instruct-2409(Mistral-Small). These mid-sized models (approximately 24–42 billion parameters) allow us to examine trade-offs compared to their larger counterparts, if any [38, 39].

#### 2.2.3 Smaller Generalist Foundation Models

Given that our downstream application may require sub-second responses, inference latency is a critical factor. Accordingly, we also investigate how the smaller model variants affect the latency–accuracy trade-off. We consider investigating the impact of smaller counterparts of the larger models we selected, including Llama-3.1-8B-Instruct(Llama-3.1-8B), Llama-3-8B-Instruct(Llama-3-8B), Qwen2.5-7B-Instruct(Qwen-7B), Ministral-8B-Instruct-2410(Ministral-8B), and Phi-4-Mini-Instruct(Phi-4-Mini)[35, 37, 39, 40].

#### 2.2.4 Reasoning Models

Given the recent interest in LLMs with advanced reasoning capabilities such as the DeepSeek R1, and keeping computational constraints in mind we opted for the distilled variants of DeepSeek R1: DeepSeek-R1-Distill-Llama-70B(DeepSeek-Llama-70B) and DeepSeek-R1-Distill-Qwen-32B(DeepSeek-Qwen-32B), their smaller counterparts DeepSeek-R1-Distill-Llama-8B(DeepSeek-Llama-8B) and DeepSeek-R1-Distill-Qwen-7B(DeepSeek-Qwen-7B). These distilled variants are the result of fine-tuning samples from the original DeepSeek-R1 model; they are said to capture the reasoning patterns of the original [41]. Additionally, we include QwQ-32B from the Qwen team, also known for its strong reasoning abilities[42].

#### 2.2.5 Domain-specific Biomedical Models

Given the biomedical focus of our research, we include three domain-specific models: BioMistral-7B, a compact model tailored for biomedical applications; Meditron-70B, which is pre-trained on a curated medical corpus and selected PubMed articles; and PMC-LLaMA-13B, trained on a large-scale corpus of 4.8 million biomedical papers and subsequently instruction-tuned using a domain-specific dataset [43–45]. These models were selected based on their strong performance on the PubMedQA benchmark and to ensure diversity in model size and training scope.[46]

### 2.3 Prompting Strategies

#### 2.3.1 Zero-Shot

Zero-shot prompting techniques have removed the need for any domain-specific training data; instead, they rely on well-crafted prompts that would act as a guideline for the model performing novel tasks [47]. The model receives a task description in the prompt, and the model would then utilize its pre-existing knowledge along with the prompt to generate predictions [48].

The effectiveness of using task-specific prompts for each model has been studied extensively in the past. In previous work, prompt design strategies were shown to benefit from including: (1) task instruction, (2) input data (e.g., abstract or title), and (3) constraints on the output format, which we adopt in constructing our zero-shot prompt [49].

Considering our classification task involves assisting the model in reviewing a particular abstract and its title, we note that this is a common practice in abstract screening for systematic review and meta-analysis studies. We review previous studies on LLM-based approaches for abstract screening and replicate the prompting strategy they propose, closely following one established method while adhering to rules outlined in prior work. [26, 49]. Hence, while constructing the prompts, we adopted a standardized approach to mimic a typical interaction between a senior researcher and a research assistant. In addition, to ensure consistency in the responses, we require that the results of the LLM be in a dictionary format {“unrelated”: 0, “relationship”: 1} or {“unrelated”: 1, “relationship”: 0} format, indicating whether each abstract met the inclusion criteria. The decision to restrict the model to only output a dictionary format was to combat the challenge posed by the LLMs free-text nature, hence in the case where the model tends to be more verbose, it largely includes a dictionary containing the final result in the response, we can then proceed to extract the verdict using custom parsing scripts. Below is the prompt used in our Zero-Shot experiment :

Zero-Shot Prompt

I seek assistance with a systematic review focused on the direct relationship between pathogens and diseases, specifically {disease}.

I’ll provide the title and abstract of a particular journal article and would appreciate an assessment for its inclusion based on the following criteria:

1. The title or abstract provides sufficient evidence of a direct relationship between the disease ({disease}) and the pathogen ({pathogen}).
2. The title or abstract investigates the Pathogen ({pathogen}) and reports evidence for the Disease ({disease}).
3. The title or abstract investigates the Disease ({disease}) and reports evi-dence for the Pathogen ({pathogen}).
4. The title or abstract states the association between the Pathogen ({pathogen}) and the Disease ({disease}), but does not focus on it.
5. The title and abstract present data or findings supporting this association.

Exclusion criteria

1. The title and abstract do not provide sufficient evidence of a direct relationship between the disease ({disease}) and the pathogen ({pathogen}).

Please provide the assessment in the following dictionary format:

{“relationship”: 1, “unrelated”: 0} if there is a relationship, or {“relationship”: 0, “unrelated”: 1} if the study should be excluded.

Note: only one value can be 1 at a time.

Title: {title}

Abstract: {abstract}

You are required to classify a journal article based solely on the given title and abstract. Do not use any external knowledge or assumptions beyond the text provided. Your decision must be strictly based on the information within the title and abstract.

Respond only in the dictionary format with no explanation. Answer:

#### 2.3.2 Few-Shot

In few-shot prompting, we provide the models with a few input-output Examples to provide an understanding of the given task, unlike zero-shot prompting, where no examples are given [49]. However, few-shot prompting requires additional tokens to include examples. The selection and composition of prompt examples can have a significant influence on how the model responds to the question. [48] In our case, we manually select examples of abstracts, belonging to the following 3 groups:

- Spurious: Abstract with a spurious relation between the disease and the pathogen term
- Not Significant: Abstract where they investigate the relation between disease and pathogen, but the pathogen didn’t show any significant results.
- Significant: Abstract where they investigate the relation between disease and pathogen, and have significant results.

Abstracts were manually labeled into one of the three groups during the dataset annotation phase. We opt for a 3-shot and 6-shot approach for this study, and with every experiment, we ensure abstracts belonging to all three groups are represented in the examples. The 6 selected examples are left out and are never used to evaluate the model.

Few-Shot Prompt

I seek assistance with a systematic review focused on the direct relationship between pathogens and diseases, specifically {disease}.

I’ll provide the title and abstract of a particular journal article and would appreciate an assessment for its inclusion based on the following criteria:

1. The title or abstract provides sufficient evidence of a direct relationship between the disease ({disease}) and the pathogen ({pathogen}).
2. The title or abstract investigates the Pathogen ({pathogen}) and reports evidence for the Disease ({disease}).
3. The title or abstract investigates the Disease ({disease}) and reports evidence for the Pathogen ({pathogen}).
4. The title or abstract states the association between the Pathogen ({pathogen}) and the Disease ({disease}), but does not focus on it.
5. The title and abstract present data or findings supporting this association.

Exclusion criteria

1. The title and abstract do not provide sufficient evidence of a direct relationship between the disease ({disease}) and the pathogen ({pathogen}).

Please provide the assessment in the following dictionary format: {“relationship”: 1, “unrelated”: 0} if there is a relationship, or {“relationship”: 0, “unrelated”: 1} if the study should be excluded.

Note: only one value can be 1 at a time.

Below are example classifications:

{examples}

Title: {title}

Abstract: {abstract}

You are required to classify a journal article based solely on the given title and abstract. Do not use any external knowledge or assumptions beyond the text provided. Your decision must be strictly based on the information within the title and abstract.

Respond only in the dictionary format with no explanation.

Answer:

#### 2.3.3 Chain-of-Thought

Chain-of-Thought (CoT) was introduced as a technique to prompt LLMs in a way that allows a step-by-step reasoning process, and this is seen to have generated more accurate results. Originally demonstrated for a given math problem, the prompt would show the reasoning process step by step, forcing the model to mimic how humans break down problems into logical intermediate steps [48]. The following is the prompt we used to enable a step-by-step reasoning process before the model makes any predictions. We intentionally omitted a few-shot exemplars in the CoT prompt to isolate the effect of verbalized reasoning :

Chain of Thought Prompt with Reasoning

Task: Classify a journal article based on whether it provides evidence of a direct relationship between {pathogen} and {disease}. Follow these steps:

1. Analyze Title:
  - Does the title explicitly mention both {pathogen} and {disease}?
  - Does it imply a relationship (e.g., “causes,” “induces,” “role in,” “association with”)?
2. Analyze Abstract:
  - Check if the abstract includes any of the following:
  - Direct claims of causation (e.g., “{pathogen} causes {disease}”).
  - Investigation of {pathogen} with findings related to {disease} (e.g., “we studied {pathogen} and observed {disease} outcomes”).
  - Study of {disease} with evidence implicating {pathogen} (e.g., “{pathogen} was identified in {disease} patients”).
  - Brief mentions of an association (e.g., “{pathogen} has been linked to {disease}”).
  - Data supporting the relationship (e.g., statistical results, experimental methods).
3. Map to Criteria:
  - C1: Direct causal claim in title/abstract.
  - C2: Abstract investigates {pathogen} and reports {disease} outcomes.
  - C3: Abstract investigates {disease} and identifies {pathogen} involvement.
  - C4: Abstract mentions association without focus.
  - C5: Abstract presents data (statistics, experiments, case studies).
4. Exclusion Check:
  - E1: No mention of both {pathogen} and {disease}, or explicitly states no relationship.
5. Decision:
  - If ANY of C1–C5 are met AND E1 is not triggered →???Include ({{“relationship”: 1, “unrelated”: 0}}).
  - If E1 applies →?Exclude ({{“relationship”: 0, “unrelated”: 1}}).

Title: {title}

Abstract: {abstract}

Reasoning:

- Title Analysis: [Mentions both? Implies relationship?]
- Abstract Findings: [Specific phrases/data related to criteria C1–C5]
- Criteria Met: [List which criteria apply]
- Exclusion Check: [Does E1 apply?]
- Final Decision: [Include/Exclude]

Answer:

### 2.4 Evaluation

Evaluating the outputs of LLMs is challenging due to their free-text nature, which often results in noisy or partially correct responses that don’t fit neatly into predefined categories. To control this, we constrained the models to respond using a custom dictionary format, and we developed custom parsing scripts for each model, which allowed us to extract only meaningful answers from the response. Any response that did not match the expected dictionary pattern was flagged as invalid and logged.

The presence of invalid outputs made it challenging to fairly evaluate model performance, so we analyzed the results in two different ways. In the strict view, every invalid response is counted as an incorrect prediction (unrelated). In the lenient view, we exclude invalid responses from the calculation entirely, looking only at the cases where the model gave a valid answer.

Once we standardized the output, we evaluated the model’s performance using a set of metrics. Specifically, we measured: (1) Avg. Query Latency (s), Elapsed time per query; (2) Balanced Accuracy, the average of sensitivity and specificity, accounting for class imbalance; (3) Precision, the accuracy of positive predictions; (4) Recall, the true positives rate; (5) Macro-F1, balancing precision and recall across classes; (6) Specificity, the true negative rate; (7) MCC (Matthews Correlation Coefficient), a balanced measure of binary classification quality even with class imbalance; and (8) Invalid Outputs, the proportion of malformed or unusable responses. To assess statistical stability, we applied bootstrapping with 2,000 resamples when computing confidence intervals for performance metrics.

To ensure consistent and deterministic outputs, all model generations were produced using greedy decoding (temperature = 0, random seed = 42). The default maximum output cap was set to 300 tokens, which is sufficient for the dictionary output plus minimal overflow. However, for models like Qwen-72B, which we noticed often truncated mid-key, we increased the cap to 1,000 tokens. Similarly, CoT runs and reasoning models were given a 1,000-token limit to ensure the full reasoning tokens are accommodated within the response.

Prompting strategies were tailored to each model’s capabilities. Generalist foundation models were tested with zero-shot, few-shot (3-shot and 6-shot), and CoT prompting. However, for reasoning models (like DeepSeek-based models and QwQ-32B), we skipped explicit CoT prompts. For domain-specific biomedical models (e.g., BioMistral-7B, PMC-LLaMA-13B), we limited evaluations to zero-shot only, since their smaller context windows made few-shot and CoT prompts unstable.

## 3 Results

To explore the potential of large language models (LLMs) in biomedical information extraction, we focus on a classification task that asks models to determine whether a given title and abstract state a direct relationship between a pathogen and a disease.

As described earlier, we evaluate predictions under two scoring policies: a lenient view, where invalid outputs are dropped, and a strict view, where such outputs are treated as negatives. The lenient approach forms the basis of our main analysis and visualisations, as it isolates classification ability from formatting robustness. However, we also report strict-view results for selected models to assess how brittle each system is to format constraints.

**Table 2.** Performance of Zero-shot techniques.

| Model | Precision | Recall | Macro F1 | Specificity | Balanced Accuracy | MCC | Invalid Outputs (%) | Avg. Query Time (s) |
| --- | --- | --- | --- | --- | --- | --- | --- | --- |
| QwQ-32B | 0.52 (0.51-0.57) | 0.79<br>(0.76-0.86) | 0.68<br>(0.66-0.72) | 0.63<br>(0.59-0.67) | 0.71<br>(0.69-0.75) | 0.39<br>(0.36-0.48) | 1.91% | 0.30 |
| DeepSeek-R1-Distill-Qwen-32B | 0.53 (0.51-0.56) | 0.95<br>(0.93-0.98) | 0.69<br>(0.67-0.72) | 0.56<br>(0.51-0.60) | 0.75<br>(0.73-0.78) | 0.50<br>(0.46-0.55) | 0.00% | 0.05 |
| Meditron-70B | 0.40 (0.39-0.42) | 0.90<br>(0.87-0.94) | 0.49<br>(0.47-0.53) | 0.29<br>(0.25-0.33) | 0.60<br>(0.57-0.63) | 0.22<br>(0.17-0.28) | 2.52% | 3.72 |
| PMC.LLaMA.13B | 0.35 (0.21-0.51) | 0.05<br>(0.03-0.08) | 0.43<br>(0.41-0.46) | 0.95<br>(0.93-0.97) | 0.50<br>(0.49-0.52) | 0.00<br>(-0.06-0.08) | 17.10% | 0.96 |
| **Qwen2.5-72b-Instruct** | 0.70 (0.67-0.76) | 0.79<br>(0.76-0.85) | **0.80**<br>**(0.78-0.83)** | 0.83<br>(0.80-0.86) | **0.81**<br>**(0.79-0.84)** | **0.60**<br>**(0.56-0.67)** | 1.11% | 3.71 |
| Phi-3.5-MoE-Instruct | 0.51 (0.49-0.54) | 0.91<br>(0.87-0.94) | 0.67<br>(0.64-0.70) | 0.54<br>(0.50-0.59) | 0.73<br>(0.70-0.75) | 0.44<br>(0.39-0.49) | 0.70% | 3.24 |
| **Mistral-Small-Instruct-2409** | 0.69 (0.65-0.72) | 0.82<br>(0.77-0.86) | **0.80**<br>**(0.77-0.82)** | 0.81<br>(0.77-0.84) | **0.81**<br>**(0.78-0.84)** | **0.61**<br>**(0.54-0.65)** | 0.00% | 0.20 |
| Llama-3.3-70B-Instruct | 0.53 (0.51-0.56) | 0.95<br>(0.93-0.98) | 0.69<br>(0.67-0.72) | 0.56<br>(0.52-0.60) | 0.75<br>(0.73-0.78) | 0.50<br>(0.46-0.54) | 0.00% | 0.52 |
| Llama-3.1-70B-Instruct | 0.53 (0.51-0.56) | 0.96<br>(0.95-0.99) | 0.70<br>(0.67-0.73) | 0.56<br>(0.52-0.60) | 0.76<br>(0.74-0.79) | 0.51<br>(0.48-0.56) | 0.00% | 0.39 |
| **Llama-3.1-Nemotron-70B-Instruct** | **0.60 (0.57-0.63)** | 0.91<br>(0.89-0.95) | **0.75**<br>**(0.73-0.78)** | 0.68<br>(0.63-0.71) | **0.80**<br>**(0.77-0.82)** | **0.56**<br>**(0.52-0.61)** | 0.00% | 0.39 |
| DeepSeek-R1-Distill-Llama-70B | 0.57 (0.54-0.60) | 0.90<br>(0.88-0.95) | 0.73<br>(0.70-0.76) | 0.64<br>(0.60-0.68) | 0.77<br>(0.75-0.80) | 0.52<br>(0.48-0.58) | 0.00% | 0.29 |
| DeepSeek-R1-Distill-Qwen-7B | 0.34 (0.35-0.35) | 1.00<br>(0.99-1.00) | 0.26<br>(0.26-0.27) | 0.00<br>(0.00-0.01) | 0.50<br>(0.50-0.50) | 0.01<br>(-0.05-0.06) | 0.00% | 0.05 |
| BioMistral-7B | 0.36 (0.36-0.37) | 0.97<br>(0.94-0.98) | 0.35<br>(0.33-0.38) | 0.10<br>(0.08-0.13) | 0.53<br>(0.52-0.55) | 0.12<br>(0.07-0.17) | 0.00% | 0.07 |
| Llama-3-8B-Instruct | 0.65 (0.61-0.70) | 0.58<br>(0.52-0.62) | 0.71<br>(0.68-0.74) | 0.84<br>(0.81-0.87) | 0.71<br>(0.67-0.73) | 0.43<br>(0.36-0.49) | 0.00% | 0.33 |
| Phi-4-Mini-Instruct | 0.48 (0.45-0.51) | 0.75<br>(0.69-0.78) | 0.63<br>(0.60-0.65) | 0.58<br>(0.53-0.61) | 0.66<br>(0.63-0.68) | 0.31<br>(0.24-0.35) | 2.72% | 0.32 |
| DeepSeek-R1-Distill-Llama-8B | 0.40 (0.39-0.41) | 0.97<br>(0.94-0.98) | 0.47<br>(0.44-0.49) | 0.23<br>(0.20-0.26) | 0.60<br>(0.58-0.62) | 0.26<br>(0.21-0.29) | 0.00% | 0.79 |
| Qwen2.5-7B-Instruct | 0.51 (0.49-0.56) | 0.63<br>(0.58-0.69) | 0.65<br>(0.62-0.68) | 0.69<br>(0.66-0.73) | 0.66<br>(0.63-0.69) | 0.31<br>(0.25-0.37) | 3.22% | 0.19 |
| Ministral-8B-Instruct-2410 | 0.43 (0.42-0.45) | 0.96<br>(0.93-0.98) | 0.55<br>(0.52-0.58) | 0.34<br>(0.31-0.38) | 0.65<br>(0.63-0.67) | 0.34<br>(0.29-0.37) | 0.00% | 0.05 |
| Llama-3.1-8B-Instruct | 0.66 (0.62-0.71) | 0.66<br>(0.60-0.70) | 0.74<br>(0.71-0.77) | 0.82<br>(0.79-0.85) | 0.74<br>(0.71-0.77) | 0.48<br>(0.42-0.54) | 0.00% | 0.15 |

### 3.1 Baseline Zero-shot Performance

To establish a performance baseline, we evaluated 19 open-weight LLMs under a zeroshot prompting condition, where models were presented only with the task description and no in-context examples.

Among all models, Qwen2.5-72B and Mistral-Small achieved the highest zero-shot performance, both with a balanced accuracy of 0.81 (Qwen2.5: 0.79–0.84; Mistral-Small: 0.78–0.84). Qwen2.5-72B scored 0.80 Macro-F1 (0.78–0.83) and 0.60 MCC (0.56–0.67), but exhibited a non-negligible invalid output rate of 1.11%. In contrast, Mistral-Small matched the top accuracy while achieving the highest MCC (0.61; 0.54–0.65) and producing 0% invalid outputs, making it one of the most robust high-performing models. These top-performing models were also among the most balanced: both paired high recall (Qwen2.5-72B: 0.79 [0.76–0.85]; Mistral-Small: 0.82 [0.77–0.86]) with solid precision (Qwen2.5-72B: 0.70 [0.67–0.76]; Mistral-Small: 0.69 [0.65–0.72]), which likely contributed to their leading MCC values and strong overall performance. However, while Qwen2.5-72B led in accuracy, it diverged substantially in classification behavior. A heatmap analysis of inter-model agreement revealed that Qwen2.5-72B shared relatively low prediction similarity with other models (Figure 1).

**Fig. 1.**
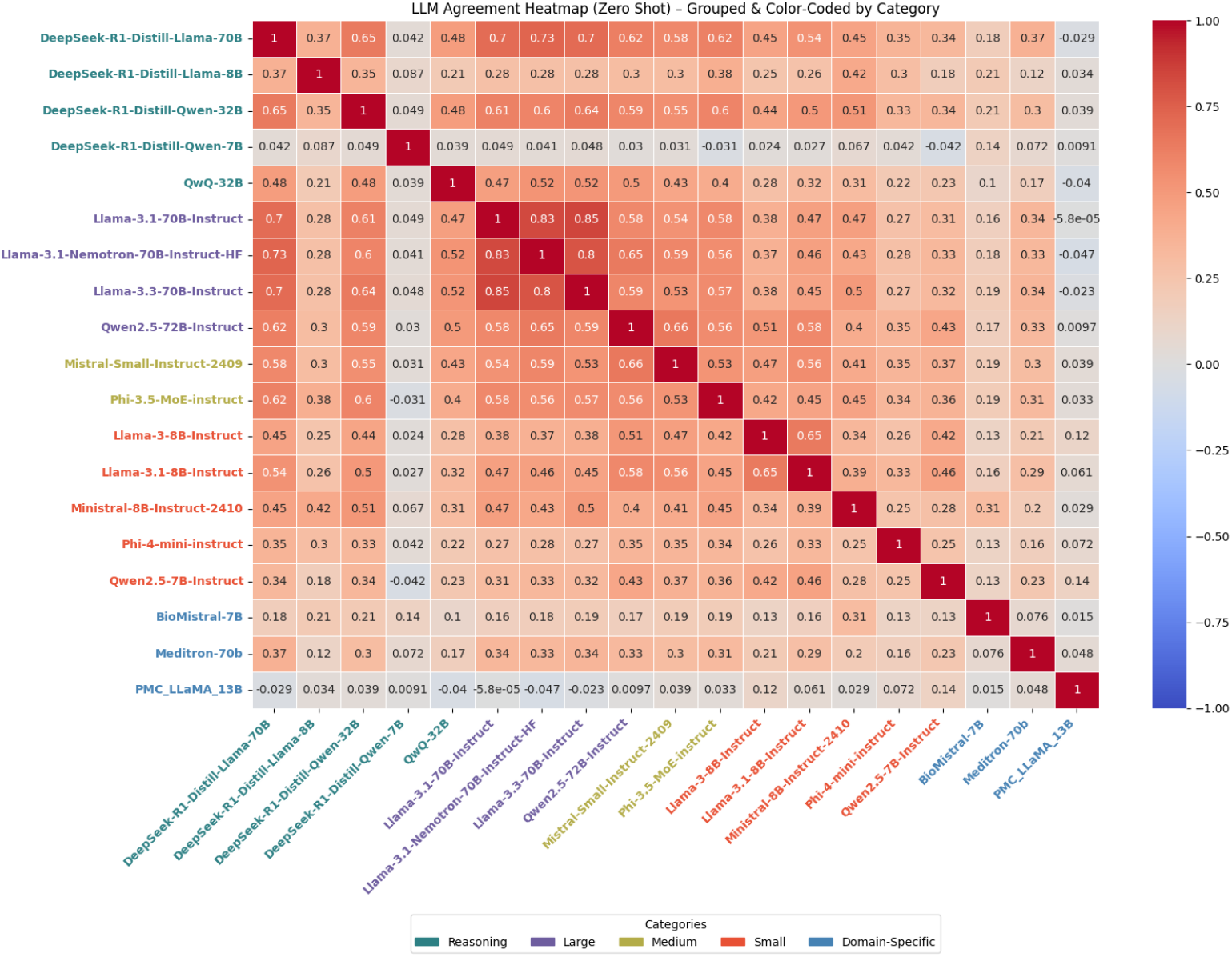
Zero-shot LLM agreement heatmap, grouped by category.

Nemotron-70B also performed competitively, with a balanced accuracy of 0.80 (0.77–0.82), Macro-F1 of 0.75 (0.73–0.78), MCC of 0.56 (0.52–0.61), and no invalid outputs (0.00%). This model, along with other Llama-family variants (e.g., Llama-3.3-70B and Llama-3.1-70B), demonstrated strong internal consistency, as evidenced by high inter-model agreement in the heatmap (Figure 1). Most generalist models in the Llama, Qwen, and Mistral families clustered near this precision–recall balance.

To test whether these observed differences were statistically meaningful, we computed paired-bootstrap (2,000×) 95% confidence intervals (CIs) on Δ-Balanced-Accuracy. The results confirm that Qwen2.5-72B trails both Nemotron-70B ((Δ = –0.041, 95% CI = [–0.064, –0.018]) and Mistral (Δ = –0.059, 95% CI = [–0.087, –0.031]), whereas Mistral-Small and Nemotron-70B form a statistical tie (Δ = +0.018, CI –0.010 … +0.046).

These results demonstrate that generalist LLMs can deliver strong zero-shot performance on structured biomedical text classification tasks. In the context of pathogen–disease relation classification, models such as Mistral-Small and the Llama variants offer a convincing combination of accuracy and robustness.

### 3.2 Effect of Prompting Strategies on Performance

Building on the established zero-shot baselines, we now examine how alternative prompting strategies, few-shot (3-shot and 6-shot) and CoT, influence model performance. These configurations test whether additional contextual examples or explicit reasoning paths improve the classification of pathogen–disease relationships.

#### 3.2.1 Few-Shot

A considerable number of models demonstrated performance degradation with the introduction of in-context examples (Figure 3). For instance, Qwen2.5-72B, despite strong zero-shot performance (Balanced Accuracy: 0.81 [0.79–0.84]; MCC: 0.60 [0.56–0.67]), experienced a dramatic increase in invalid outputs to 10.76% under 3-shot prompting (Figure 4). This sharp rise in invalid completions undermines the apparent performance, indicating an overestimation when only valid outputs are considered. Similarly, Llama-3.1-Nemotron saw a decline in both balanced accuracy and MCC as prompting depth increased. Its MCC dropped from 0.56 (0.52–0.61) (zero-shot) to 0.40 (0.37–0.46) (6-shot), primarily driven by a drop in precision from 0.60 (0.57–0.63) to 0.47 (0.47–0.51). Balanced accuracy followed suit, declining from 0.80 (0.77–0.82) to 0.70 (0.68–0.73). Other models, including Phi-3.5-MoE and Llama-3.1-8B, showed similar regressions: Phi-3.5-MoE’s MCC fell from 0.44 (0.39–0.49) to 0.28 (0.22–0.33), and invalid completions rose to 3.62%. Meanwhile, Llama-3.1-8B’s MCC plummeted to 0.18 (0.12–0.23), with invalid outputs increasing to 3.52%.

**Fig. 2.**
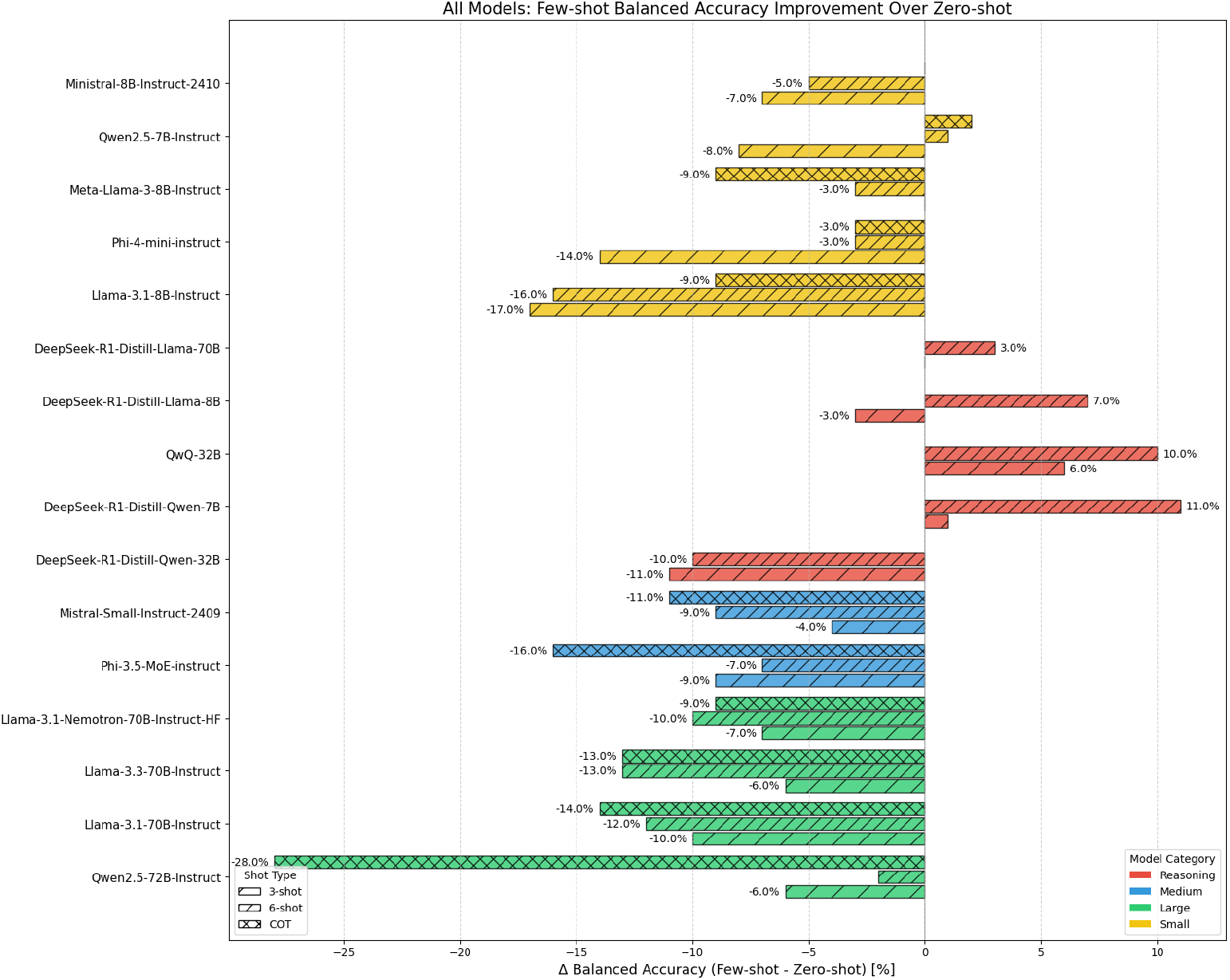
Few-shot and CoT Balanced Accuracy Improvement per Model.

**Fig. 3.**
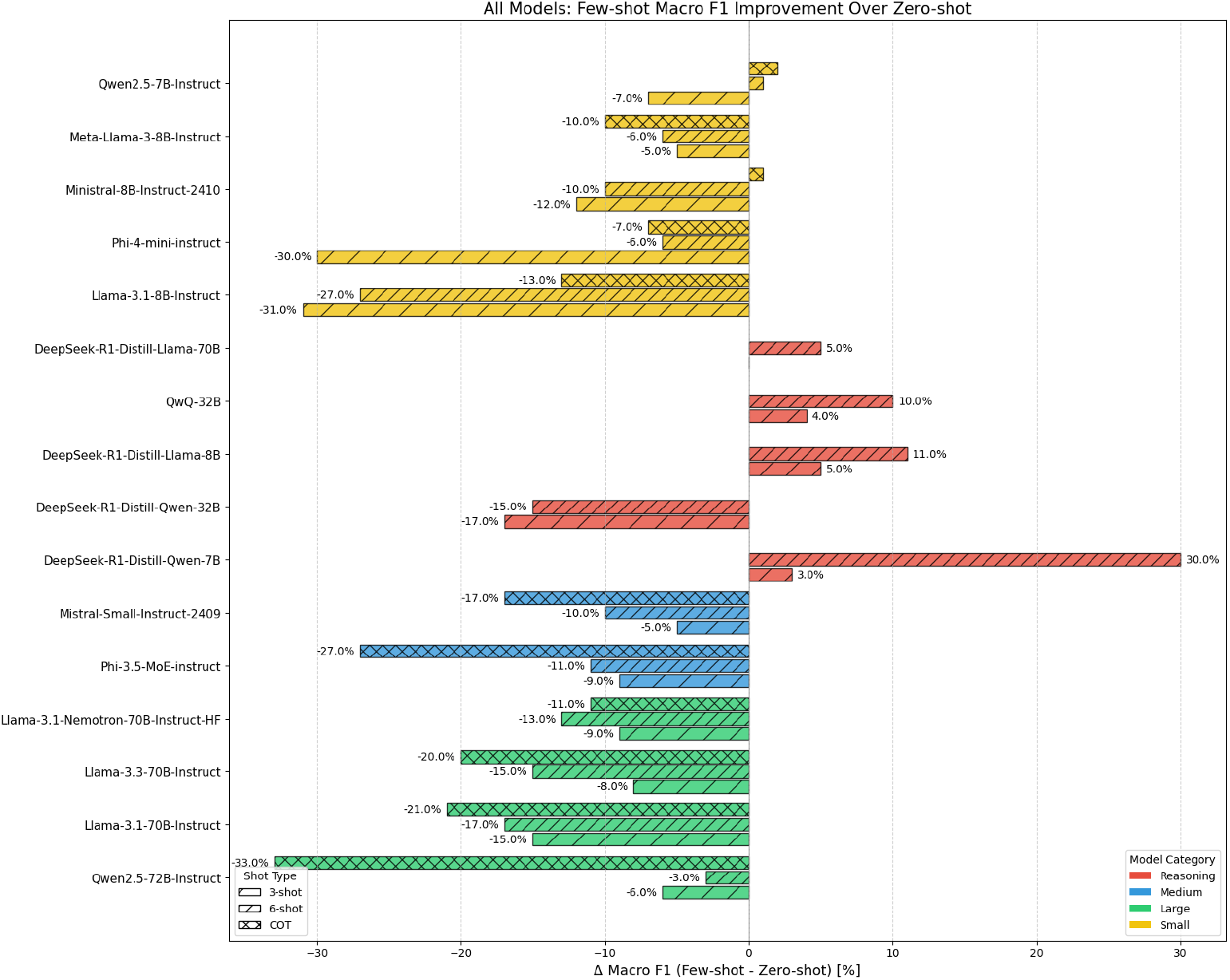
Few-shot and CoT Macro F1 Improvement per Model.

**Fig. 4.**
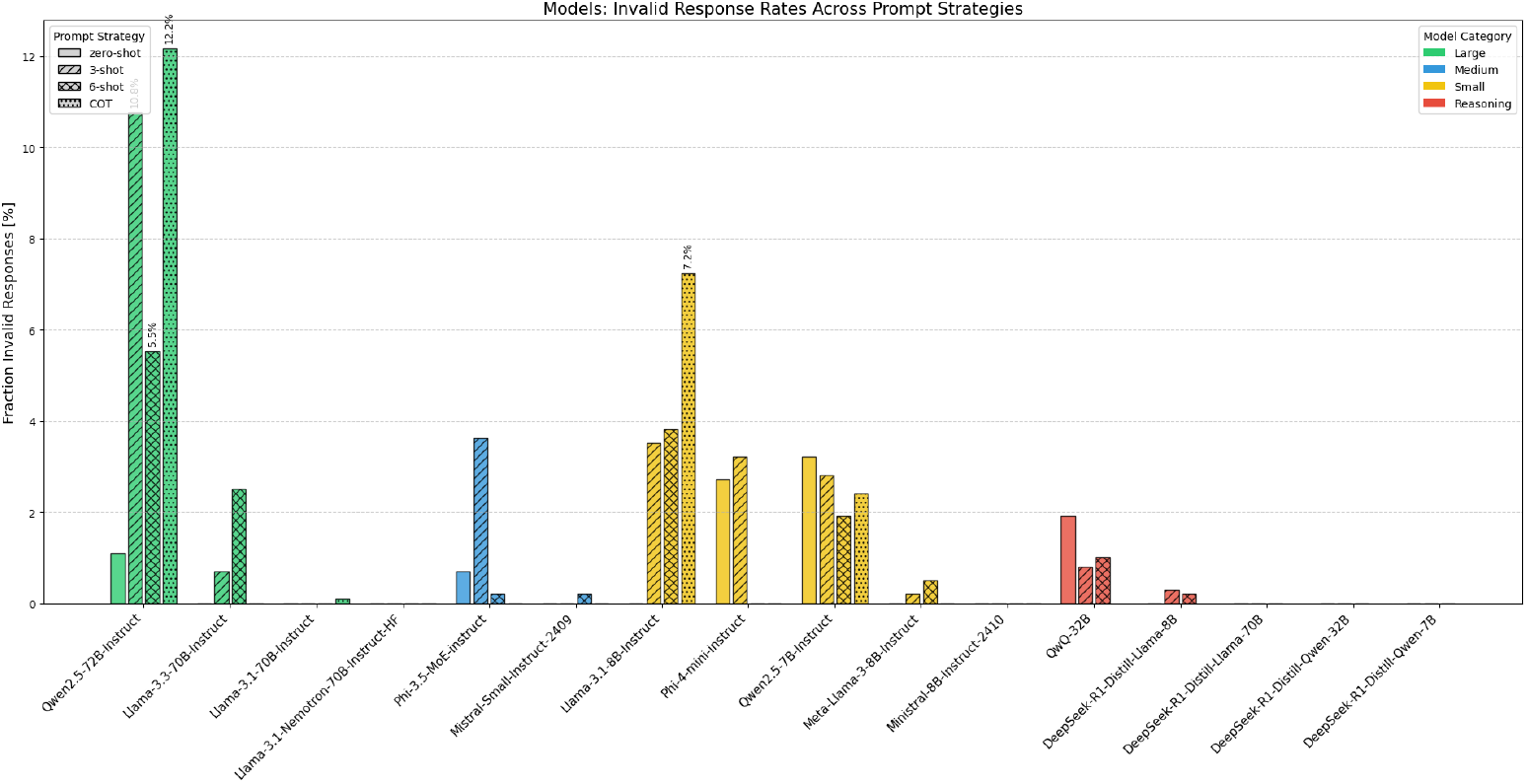
Invalid Output Rate per Model.

A distinct case is DeepSeek-Qwen-32B, which, despite achieving extremely high recall at 3-shot (0.98 [0.96–0.99]), exhibited low specificity (0.30 [0.25–0.33]) and modest MCC (0.33 [0.29–0.36]). Although its balanced accuracy appeared stable at 0.64 (0.62–0.66), the imbalance between precision (0.42 [0.42–0.45]) and recall resulted in unreliable overall predictions.

In contrast, reasoning models demonstrated improved or consistently strong performance as prompt depth increased. DeepSeek-Llama-70B was one such example, showing steady gains in all core metrics: Balanced Accuracy improved from 0.77 (0.75–0.80) to 0.80 (0.78–0.83), Macro F1 rose from 0.73 (0.70–0.76) to 0.78 (0.76–0.81), and MCC increased from 0.52 (0.48–0.58) to 0.58 (0.54–0.64). Importantly, the model maintained a 0.00% invalid output rate across all prompting conditions.

QwQ-32B also benefited substantially from few-shot prompting (Figure 3). It demonstrated improvements from zero-shot to 6-shot: Balanced Accuracy increased from 0.71 (0.69–0.75) to 0.81 (0.79–0.84), Macro F1 from 0.68 (0.66–0.72) to 0.78 (0.76–0.81), and MCC from 0.39 (0.36–0.48) to 0.59 (0.55–0.64). Precision improved from 0.52 (0.51–0.57) to 0.64 (0.61–0.68), while recall remained consistently high (from 0.79 [0.76–0.86] to 0.88 [0.86–0.92]). Although invalid outputs ranged from 0.80% (3-shot) to 1.91% (zero-shot), the overall metric gains marked this model as among the most responsive to few-shot prompting.

**Table 3.**
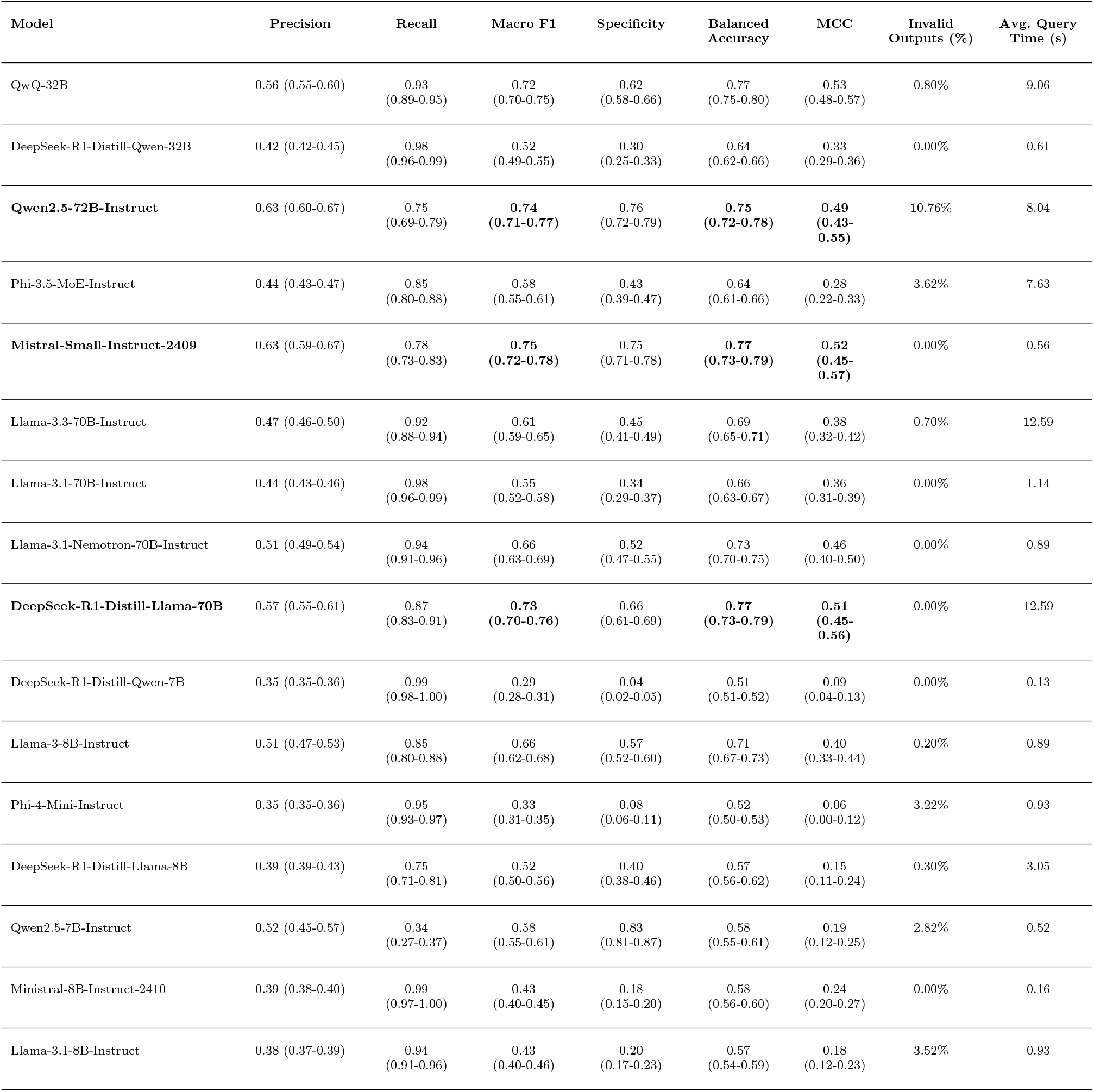
Performance of LLMs 3-shot.

A subset of models exhibited inconsistent performance trends across prompt depths. Phi-4-Mini, for instance, experienced an initial performance decline at 3-shot but showed partial recovery at 6-shot, with gains in precision and macro F1 relative to the 3-shot setting. However, these gains remained insufficient to surpass its original zero-shot performance.

Few-shot prompting did not lead to consistent improvements across models; in most cases, performance declined, and invalid outputs generally increased with prompt depth(Figure 2). Reasoning models, such as QwQ-32B and DeepSeek-Llama-70B, were notable exceptions, showing reliable gains.

#### 3.2.2 Chain-of-Thought

The application of CoT prompting generally resulted in pronounced performance degradation relative to both zero-shot and few-shot prompting strategies (Figure 2).

Qwen-72B, which achieved strong baseline performance in the zero-shot setting (Balanced Accuracy: 0.81 [0.79–0.84]; MCC: 0.60 [0.56–0.67]), suffered the most substantial decline under CoT prompting. Balanced accuracy fell sharply to 0.53 (0.50–0.57), and MCC collapsed to 0.07 (0.00–0.14). Compounding this degradation was a more than tenfold increase in invalid outputs, rising from 1.11% to 12.17%, severely undermining the model’s reliability. These declines were more severe than those observed under 3-shot prompting, where balanced accuracy was 0.75 (0.72–0.78), MCC was 0.49 (0.43–0.55), and invalid outputs were already elevated at 10.76

Llama-3.1-70B, another model with relatively high zero-shot performance, also experienced significant deterioration under CoT. Balanced accuracy dropped to 0.62 (0.60–0.64) and MCC to 0.31 (0.26–0.33), marking reductions from both its zero-shot and few-shot values. Although invalid output rates remained low (0.10%), precision and specificity declined notably, with the latter falling from 0.56 to 0.26. Recall, however, remained extremely high (≥ 99%) under all prompting strategies, suggesting that CoT shifted the model toward high-recall, low-specificity behavior, a pattern recurrent across other models as well.

In many cases, CoT prompting increased recall while compromising other facets of performance. Phi-3.5-MoE, for example, improved recall from 0.91 (0.87–0.94) to 0.98 (0.96–1.00) under CoT, but this came at the expense of precision and specificity, which dropped to 0.38 and 0.15, respectively. The result was a sharp decline in MCC from 0.44 (0.39–0.49) to 0.20 (0.15–0.24), below its 3-shot level (0.28).

By contrast, Ministral-8B maintained stable and balanced performance across all prompting strategies. Under CoT, recall slightly declined (from 0.96 to 0.94), but this was accompanied by consistent precision (0.44), marginal improvement in specificity (0.34 →?0.36), and negligible MCC change (0.34 →?0.33). Importantly, the model preserved a 0.00% invalid output rate across zero-shot, few-shot, and CoT.

Other models, such as Llama-3.1-8B, Phi-4-Mini, and Qwen-7B, demonstrated moderate declines under CoT prompting. For Llama-3.1-8B, recall decreased to 0.75 (0.70–0.81), precision to 0.46 (0.44–0.49), and MCC to 0.28 (0.22–0.34), with invalid completions rising to 7.24%. Phi-4-Mini followed a similar trajectory, with recall at 0.87 (0.82–0.90), precision at 0.43 (0.41–0.45), and MCC at 0.27 (0.21–0.33), though it avoided invalid completions entirely (0.00%). Qwen-7B recorded a moderate invalid rate (2.41%) and an MCC of 0.35 (0.32–0.45).

Specificity emerges as an important factor to differentiate CoT from few-shot prompting. Under CoT, it dropped steeply in models that had otherwise maintained balanced true positive and true negative rates. As seen with Phi-3.5-MoE and Llama-3.1-70B, specificity reductions (e.g., 0.54 →0.15 and 0.56 →0.26, respectively) led to a precision–recall imbalance that skewed predictions toward the positive class, degrading classification reliability.

Compared to zero-shot and few-shot prompting, CoT strategies consistently underperformed across a range of models and introduced greater output invalidity. While few-shot prompting occasionally led to moderate declines, as observed in Qwen2.5-72B and Nemotron-70B, CoT prompting worsened these effects. Qwen-72B’s MCC declined by 0.11 under 3-shot prompting but by 0.53 under CoT, along with a substantial rise in invalid completions (Figure 4). Even typically robust models like Llama-3.1-70B suffered an average MCC reduction of 0.19, emphasizing the destabilizing effect of CoT in this case. These degradations were frequently driven by severe imbalances in recall and specificity. Overall, CoT prompting introduced more pronounced trade-offs than either zero-shot or few-shot prompting, suggesting it may be unsuitable for our specific classification task.

**Table 4.** Performance of LLMs using 6-shot prompting.

| Model | Precision | Recall | Macro F1 | Specificity | Balanced Accuracy | MCC | Invalid Outputs (%) | Avg. Query Time (s) |
| --- | --- | --- | --- | --- | --- | --- | --- | --- |
| **QwQ-32B** | 0.64 (0.61-0.68) | 0.88<br>(0.86-0.92) | **0.78**<br>**(0.76-0.81)** | 0.74<br>(0.70-0.77) | **0.81**<br>**(0.79-0.84)** | **0.59**<br>**(0.55-0.64)** | 1.01% | 0.79 |
| DeepSeek-R1-Distill-Qwen-32B | 0.43 (0.43-0.46) | 0.97<br>(0.95-0.99) | 0.54<br>(0.52-0.58) | 0.33<br>(0.29-0.37) | 0.65<br>(0.63-0.67) | 0.34<br>(0.31-0.38) | 0.00% | 1.03 |
| **Qwen2.5-72B-Instruct** | 0.67 (0.64-0.72) | 0.77<br>(0.72-0.82) | **0.77**<br>**(0.75-0.81)** | 0.80<br>(0.77-0.84) | **0.79**<br>**(0.76-0.82)** | **0.55**<br>**(0.51-0.62)** | 5.53% | 13.77 |
| Phi-3.5-MoE-Instruct | 0.44 (0.43-0.46) | 0.94<br>(0.92-0.97) | 0.56<br>(0.53-0.59) | 0.37<br>(0.32-0.40) | 0.66<br>(0.63-0.68) | 0.34<br>(0.29-0.38) | 0.20% | 11.72 |
| Mistral-Small-Instruct-2409 | 0.55 (0.53-0.59) | 0.75<br>(0.71-0.80) | 0.70<br>(0.67-0.73) | 0.68<br>(0.64-0.72) | 0.72<br>(0.69-0.75) | 0.41<br>(0.36-0.47) | 0.20% | 1.12 |
| Llama-3.3-70B-Instruct | 0.42 (0.41-0.45) | 0.90<br>(0.86-0.93) | 0.54<br>(0.51-0.57) | 0.35<br>(0.31-0.39) | 0.62<br>(0.60-0.65) | 0.26<br>(0.21-0.32) | 2.52% | 13.15 |
| Llama-3.1-70B-Instruct | 0.42 (0.42-0.45) | 0.98<br>(0.96-0.99) | 0.53<br>(0.51-0.56) | 0.31<br>(0.27-0.35) | 0.64<br>(0.62-0.66) | 0.33<br>(0.30-0.37) | 0.00% | 1.91 |
| Llama-3.1-Nemotron-70B-Instruct | 0.47 (0.47-0.51) | 0.94<br>(0.92-0.97) | 0.62<br>(0.60-0.66) | 0.45<br>(0.42-0.50) | 0.70<br>(0.68-0.73) | 0.40<br>(0.37-0.46) | 0.00% | 1.91 |
| **DeepSeek-R1-Distill-Llama-70B** | 0.64 (0.62-0.68) | 0.87<br>(0.83-0.91) | **0.78**<br>**(0.76-0.81)** | 0.74<br>(0.71-0.78) | **0.80**<br>**(0.78-0.83)** | **0.58**<br>**(0.54-0.64)** | 0.00% | 19.50 |
| DeepSeek-R1-Distill-Qwen-7B | 0.42 (0.40-0.45) | 0.75<br>(0.71-0.80) | 0.56<br>(0.54-0.60) | 0.47<br>(0.43-0.51) | 0.61<br>(0.58-0.65) | 0.21<br>(0.16-0.28) | 0.00% | 0.08 |
| Llama-3-8B-Instruct | 0.50 (0.47-0.52) | 0.76<br>(0.71-0.80) | 0.65<br>(0.62-0.68) | 0.60<br>(0.57-0.64) | 0.68<br>(0.65-0.71) | 0.34<br>(0.28-0.40) | 0.50% | 1.45 |
| Phi-4-Mini-Instruct | 0.43 (0.41-0.46) | 0.80<br>(0.77-0.85) | 0.57<br>(0.55-0.61) | 0.45<br>(0.42-0.50) | 0.63<br>(0.61-0.66) | 0.25<br>(0.21-0.32) | 0.00% | 0.47 |
| DeepSeek-R1-Distill-Llama-8B | 0.45 (0.43-0.46) | 0.94<br>(0.91-0.96) | 0.58<br>(0.55-0.61) | 0.40<br>(0.36-0.44) | 0.67<br>(0.65-0.69) | 0.35<br>(0.31-0.40) | 0.20% | 1.04 |
| Qwen2.5-7B-Instruct | 0.53 (0.49-0.56) | 0.64<br>(0.59-0.69) | 0.66<br>(0.63-0.69) | 0.70<br>(0.66-0.74) | 0.67<br>(0.64-0.70) | 0.33<br>(0.27-0.39) | 1.91% | 0.79 |
| Ministral-8B-Instruct-2410 | 0.39 (0.38-0.40) | 0.99<br>(0.97-1.00) | 0.45<br>(0.42-0.48) | 0.21<br>(0.18-0.25) | 0.60<br>(0.58-0.62) | 0.26<br>(0.23-0.30) | 0.00% | 0.26 |
| Llama-3.1-8B-Instruct | 0.38 (0.37-0.40) | 0.89<br>(0.85-0.92) | 0.47<br>(0.45-0.51) | 0.27<br>(0.25-0.31) | 0.58<br>(0.56-0.61) | 0.18<br>(0.13-0.24) | 3.82% | 1.45 |

### 3.3 Performance Across Model Sizes

Understanding how model scale influences performance is important for selecting the most appropriate LLM in terms of both accuracy and efficiency. In this section, we analyze models of three different scales: small, mid-sized, and large.

The larger models, typically around 70 billion parameters, such as Qwen-72B and Llama-3.1-Nemotron-70B, achieve the highest overall performance on average, with balanced accuracy reaching approximately 0.80–0.81 and macro-F1 scores between 0.75 and 0.80. Mid-size models, in the 24–32B range, often match or exceed their larger counterparts in balanced metrics. Mistral-Small, for example, achieves a macro-F1 of 0.80 (0.77–0.82), with precision of 0.69 (0.65–0.72) and recall of 0.82 (0.77–0.86), and an MCC of 0.61 (0.54–0.65), performing comparably to the Llama-70B variants. In contrast, Phi-3.5-MoE (24B) emphasizes recall (0.91 [0.87–0.94]) at the cost of precision (0.51 [0.49–0.54]), leading to a lower MCC of 0.44 (0.39–0.49).

Smaller models, such as Llama-3.1-8B, Llama-3-8B, and Qwen-7B, exhibit more varied performance. Most small Llama variants cluster around macro-F1 scores of 0.71–0.74 and MCCs in the 0.43–0.48 range, comparable to some of their 70B siblings but still trailing mid-size models like Mistral-Small. Qwen-7B shows lower overall performance, with a macro-F1 of 0.65 (0.62–0.68) and MCC of 0.31 (0.25–0.37), falling short of Qwen-72B. Ministral-8B, despite being smaller, performs modestly with a macro-F1 of 0.55 (0.52–0.58) and MCC of 0.34 (0.29–0.37).

We observe meaningful differences across scales, but improvements are not strictly linear. It is also worth noting that the number of mid-sized models in the evaluation is relatively small, which limits the strength of any broad generalizations about this category.

### 3.4 Domain-Specific, Reasoning vs. Generalist Models

Domain-specific models, which were restricted to zero-shot due to context window limitations, exhibited polarized behaviors. BioMistral-7B and Meditron-70B emphasized recall (e.g., BioMistral recall: 0.97 0.94–0.98]), but suffered from extremely low specificity (as low as 0.10 [0.08–0.13]), limiting their overall balanced accuracy and MCC. In contrast, PMC-LLaMA-13B reversed this trend, achieving high specificity of 0.95 (0.93–0.97) but near-zero recall at 0.05 (0.03–0.08), resulting in a balanced accuracy of 0.50 (0.49–0.52), MCC of 0.00 (–0.06–0.08), and a high invalid output rate of 17.10

Interestingly, while most performance assessments were focused on zero-shot due to instability in few-shot prompts for domain-specific models, reasoning models demonstrated noticeable gains when evaluated under few-shot conditions. For instance, QwQ-32B improved from 0.68 (0.66–0.72) to 0.78 (0.76–0.81) in macro-F1, and from 0.39 (0.36–0.48) to 0.59 (0.55–0.64) in MCC under 6-shot prompting. This suggests that, unlike domain-tuned or generalist models, reasoning models can employ few-shot learning effectively to refine predictions.

Overall, our trends show generalist models performing the best, particularly in zero-shot settings, but it is important to acknowledge that this comparison may not be entirely fair; many domain-specific models are built on older architectures and constrained by limited context windows. Moreover, even as other models struggled with few-shot prompting, reasoning models consistently benefited from it.

**Table 5.** Performance of LLMs using CoT prompting.

| Model | Precision | Recall | Macro F1 | Specificity | Balanced Accuracy | MCC | Invalid Outputs (%) | Avg. Query Time (s) |
| --- | --- | --- | --- | --- | --- | --- | --- | --- |
| Ministral-8B-Instruct-2410 | 0.44 (0.42-0.45) | 0.94<br>(0.91-0.97) | 0.56<br>(0.52-0.58) | 0.36<br>(0.31-0.40) | 0.65<br>(0.62-0.67) | 0.33<br>(0.27-0.37) | 0.00% | 1.05 |
| **Qwen2.5-72B-Instruct** | 0.37 (0.35-0.39) | 0.73<br>(0.67-0.78) | **0.47**<br>**(0.44-0.51)** | 0.34<br>(0.30-0.38) | **0.53**<br>**(0.50-0.57)** | **0.07**<br>**(0.00-0.14)** | 12.17% | 12.24 |
| Phi-3.5-MoE-Instruct | 0.38 (0.37-0.39) | 0.98<br>(0.96-1.00) | 0.40<br>(0.37-0.43) | 0.15<br>(0.11-0.17) | 0.57<br>(0.55-0.58) | 0.20<br>(0.15-0.24) | 0.00% | 11.95 |
| Llama-3.1-70B-Instruct | 0.41 (0.40-0.42) | 0.99<br>(0.97-1.00) | 0.49<br>(0.45-0.52) | 0.26<br>(0.21-0.28) | 0.62<br>(0.60-0.64) | 0.31<br>(0.26-0.33) | 0.10% | 3.49 |
| Phi-4-Mini-Instruct | 0.43 (0.41-0.45) | 0.87<br>(0.82-0.90) | 0.56<br>(0.53-0.59) | 0.40<br>(0.35-0.44) | 0.63<br>(0.60-0.66) | 0.27<br>(0.21-0.33) | 0.00% | 1.12 |
| **Mistral-Small-Instruct-2409** | 0.48 (0.46-0.51) | 0.92<br>(0.90-0.96) | **0.63**<br>**(0.60-0.66)** | 0.48<br>(0.42-0.51) | **0.70**<br>**(0.67-0.73)** | **0.39**<br>**(0.36-0.45)** | 0.00% | 0.20 |
| Llama-3.3-70B-Instruct | 0.41 (0.40-0.42) | 0.98<br>(0.96-0.99) | 0.49<br>(0.46-0.52) | 0.26<br>(0.21-0.28) | 0.62<br>(0.59-0.63) | 0.29<br>(0.25-0.33) | 0.00% | 5.62 |
| Llama-3-8B-Instruct | 0.46 (0.43-0.51) | 0.59<br>(0.53-0.65) | 0.61<br>(0.57-0.64) | 0.65<br>(0.60-0.68) | 0.62<br>(0.58-0.65) | 0.22<br>(0.15-0.29) | 0.00% | 2.19 |
| Qwen2.5-7B-Instruct | 0.54 (0.52-0.61) | 0.65<br>(0.62-0.73) | 0.67<br>(0.65-0.72) | 0.71<br>(0.69-0.76) | 0.68<br>(0.66-0.73) | 0.35<br>(0.32-0.45) | 2.41% | 2.13 |
| **Llama-3.1-Nemotron-70B-Instruct** | 0.49 (0.47-0.51) | 0.94<br>(0.92-0.97) | **0.64**<br>**(0.61-0.67)** | 0.48<br>(0.43-0.52) | **0.71**<br>**(0.68-0.74)** | **0.42**<br>**(0.38-0.47)** | 0.00% | 5.47 |
| Llama-3.1-8B-Instruct | 0.46 (0.44-0.49) | 0.75<br>(0.70-0.81) | 0.61<br>(0.58-0.64) | 0.54<br>(0.49-0.58) | 0.65<br>(0.61-0.68) | 0.28<br>(0.22-0.34) | 7.24% | 3.13 |

### 3.5 Latency–accuracy trade-offs across prompting strategies

Here we address the question “What latency–accuracy trade-offs emerge across models and prompt types?”, in a dedicated analysis across all prompting strategies, summarized visually in Figure 5 (latency vs. Macro-F1 scatter plot). This plot reveals a clear trade-off frontier: zero-shot prompting lies closest to the best possible curve, delivering strong accuracy with the lowest latency. For example, Mistral-Small achieves Macro-F1 scores around 0.80 in under 0.3 seconds, making it highly efficient and effective at the same time. In contrast, few-shot prompting (3- and 6-shot) in most cases increases latency by 3–5×, yet rarely improves accuracy and often degrades, especially for large models like Llama-3 70B parameter models, where Macro-F1 drops. CoT prompts further increase latency costs, pushing inference times while simultaneously lowering accuracy in most cases. The scatter plot illustrates that many CoT and fewshot points shift rightward (slower) and downward (less accurate) compared to their zero-shot baselines.

**Fig. 5.**
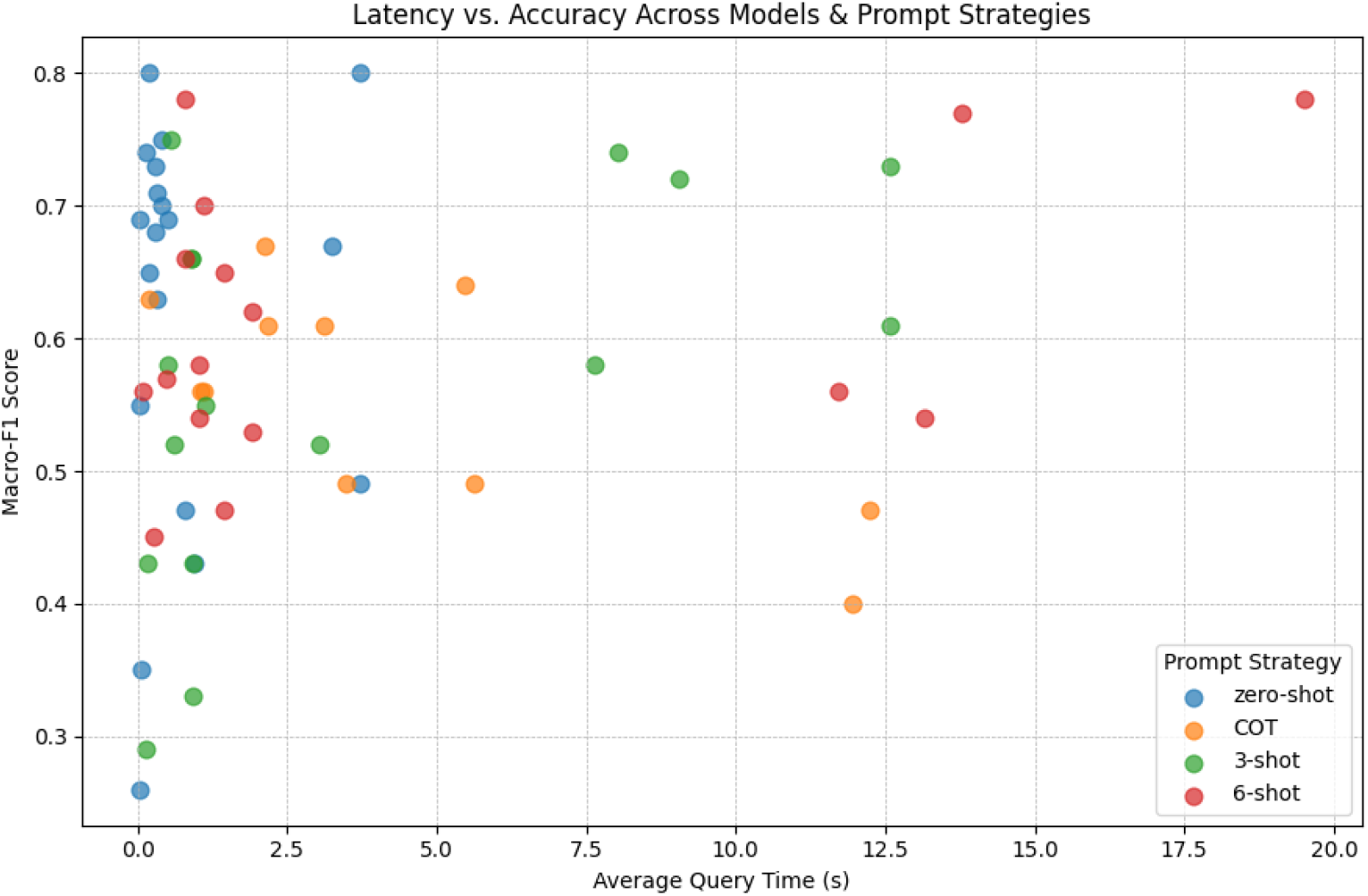
Latency vs. accuracy (Macro-F1) across models and prompt strategies.

### 3.6 Performance Robustness in Strict vs. Lenient Evaluation

To evaluate the effect of stricter output parsing criteria on our model comparisons, we reran the zero-shot evaluation using a strict grading policy, where every invalid or malformed answer is treated as an incorrect prediction. As shown in Table 6, models with invalid outputs (e.g., PMC-LLaMA-13B at 17.1%, Qwen-2.5-7B at 3.2%) saw balanced accuracy reductions of up to 1 point, though the top-line model ranking remained unchanged. The same stability held across 3-shot prompting as well, where even models with *>* 10% invalid outputs (e.g., Qwen-72B) saw ≤ 1 pp drop in balanced accuracy under the strict view (Table 7). This highlights that the performance differences we captured under the lenient view remain largely preserved in the zero-shot setting of the strict view, hence concluding for our given case, the choice of evaluation policy, whether dropping invalid outputs or treating them as incorrect, has only a minor effect on overall rankings in the zero-shot setting.

**Table 6.** Comparison of strict vs lenient evaluation in the zero-shot setting.

| Model | Prompt Strategy | Balanced Acc. (Lenient) | Balanced Acc. (Strict) | $\Delta$ BA (pp) | Invalid Outputs (%) |
| --- | --- | --- | --- | --- | --- |
| QwQ-32B | zero-shot | 0.71 | 0.70 | -1.0 | 1.91% |
| Qwen2.5-7B-Instruct | zero-shot | 0.66 | 0.65 | -1.0 | 3.22% |
| Meditron-70B | zero-shot | 0.60 | 0.59 | -1.0 | 2.52% |
| Qwen2.5-72B-Instruct | zero-shot | 0.81 | 0.80 | -1.0 | 1.11% |
| PMC-LLaMA-13B | zero-shot | 0.66 | 0.66 | 0.0 | 2.72% |

**Table 7.**
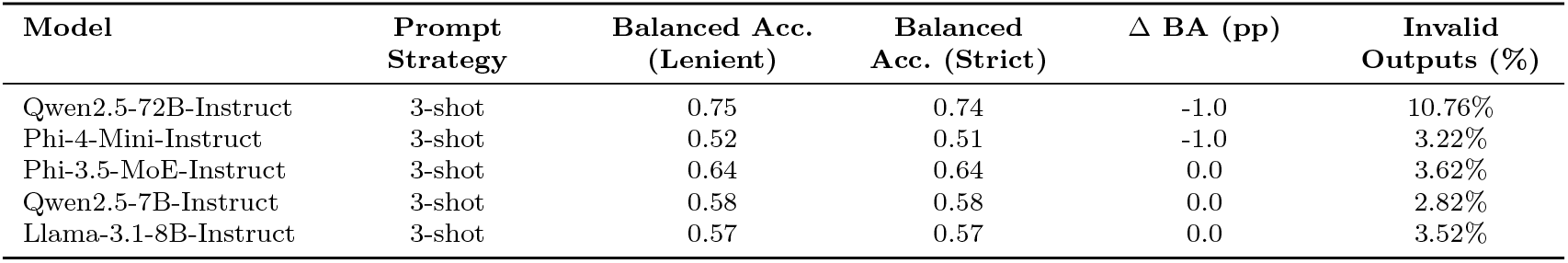
Comparison of strict vs lenient evaluation in the 3-shot prompt-strategy setting.

| Model | Prompt Strategy | Balanced Acc. (Lenient) | Balanced Acc. (Strict) | $\Delta$ BA (pp) | Invalid Outputs (%) |
| --- | --- | --- | --- | --- | --- |
| Qwen2.5-72B-Instruct | 3-shot | 0.75 | 0.74 | -1.0 | 10.76% |
| Phi-4-Mini-Instruct | 3-shot | 0.52 | 0.51 | -1.0 | 3.22% |
| Phi-3.5-MoE-Instruct | 3-shot | 0.64 | 0.64 | 0.0 | 3.62% |
| Qwen2.5-7B-Instruct | 3-shot | 0.58 | 0.58 | 0.0 | 2.82% |
| Llama-3.1-8B-Instruct | 3-shot | 0.57 | 0.57 | 0.0 | 3.52% |

## 4 Discussion

This study began with a rigorous exploration of the capabilities of LLMs in the classification of the existence of pathogen-disease relationships, given the title and abstract of a biomedical article. Our goal was to evaluate the efficiency and accuracy of multiple open-source foundational LLMs, which are of varying sizes ranging from 7B to 70B parameters, and establish their baseline capabilities. Following this, we intended to evaluate how different prompting techniques impacted the performance and efficiency.

Our findings are broadly consistent with the notion that, under certain conditions, larger models can offer performance advantages. That said, we did not observe a straightforward relationship between size and improvement. In our limited set of models, the step from roughly 7 billion to 13–24 billion parameters appeared to yield the most notable gains in balanced accuracy and macro-F1. In contrast, the increase from 33 billion to 70 billion parameters resulted in only modest or sometimes ambiguous improvements. Given the small and heterogeneous set of models evaluated, we do not suggest that this pattern is generalizable; rather, the results point to a more nuanced landscape, where architectural choices and other factors may play a significant role alongside model scale.

That said, size alone doesn’t guarantee reliability. Some of the largest models, such as Qwen-72B, scored very well on metrics like balanced accuracy and macro-F1, but these figures can be misleading. The model produced a high rate of invalid outputs, meaning many of its predictions could not be translated into predefined categories. As a result, its “high” scores are based on a reduced set of valid responses, potentially inflating its apparent performance. It is hence important to factor in output validity and robustness when evaluating the effectiveness of the model, not just raw accuracy.

Our error logs reveal three recurring failure modes behind these invalid outputs. (i) Instruction drift: models sometimes ignore the constrained “relationship”: 1, “unrelated”: 0 schema and emit a free-text explanation. (ii) Code hallucination, especially in Qwen-72B under CoT prompting, where the model prints a Python function instead of a JSON-like dictionary. (iii) Parser mismatch: our regex-based extractor assumes the final dictionary in the output is the true answer. However, if the model echoes part of the prompt, including a dictionary, and then gets truncated mid-generation, the parser mistakenly captures the wrong dictionary. A promising fix would be to introduce a two-stage pipeline: the primary model generates a response, and a lightweight verifier LLM then classifies the output into expected categories.

Our experiments revealed a counterintuitive trend: increasing the number of shots(examples) often degraded performance across generalist models of varying sizes. We observed declines in precision, balanced accuracy, and the validity of the output as more context was added. These findings align with some prior research[30], although other studies suggest that additional context can enhance performance[21, 49]. The notion that excessive context can degrade outcomes contradicts the intuitive assumption that additional examples should enhance task understanding.

One possible explanation is that the addition of context through examples introduces greater complexity, which may dilute the model’s ability to focus on the core task. In models with limited context windows, excessive prompting may push relevant parts of the query further away from the model’s attention. It should also be noted, however, that our experiments did not control for possible confounding factors such as the quality of the examples themselves. As a result, we cannot confirm whether the observed pattern would persist under more tightly controlled conditions. These findings point to potential limitations and bring about the need for further investigation into prompt length and example quality.

However, reasoning models like QwQ-32B and distilled Deepseek variants defied this general trend, showing improved performance with more context. This difference likely reflects their inherent support for self-verification and structured reasoning principles closely aligned with CoT strategies. Our attempts to manually design CoT prompts, however, led to degraded performance. We believe the limits we observed likely stem from the deliberately simplistic, example-free prompting strategy we chose. By keeping prompts brief and omitting exemplars, we underutilized established CoT techniques. Future work should ideally explore richer, exemplar-based CoT prompts. As a result, these setbacks may not indicate an inherent limitation of the technique or the model itself. A valuable direction for future work would be to analyze the reasoning patterns exhibited by high-performing reasoning models like DeepSeek-R1 distills and QwQ-32B, particularly in how they structure their thinking steps. Studying these model-generated chains of thought, one could better understand the reasoning strategies they use and adapt from these structures to improve the design of our prompts.

To assess whether domain-specific pretraining on biomedical or clinical corpora would provide an advantage for this classification task. We hence evaluated BioMistral-7B, PMC-LLaMA-13B and Meditron-70B; however, our evaluations were restricted solely to zero-shot mode because their short context windows made few-shot or CoT prompting impractical. BioMistral-7B was fast with strong recall, but PMC-LLaMA-13B often produced malformed outputs, and Meditron-70B struggled to follow reasoning cues, so none outperformed the generalist baselines. Future work should revisit this analysis with newer biomedical LLMs that pair larger context windows with modern instruction-tuned architectures.

Turning to the dataset, models performed well on abstracts with a clear structure, particularly those that explicitly stated their findings, such as ‘We investigated X and found Y. They also performed well when a significant relationship between a pathogen and disease was explicitly stated. However, performance dropped in cases where the relationship was weakly defined, negative, or buried in complex subclauses. Abstracts with multiple pathogen–disease pairs or highly condensed biomedical language caused further confusion, often leading the models to hallucinate or incorrectly pair terms based on proximity rather than context.

Latency plays a critical role in model selection, especially when LLMs are deployed in time-sensitive downstream tasks. Our evaluation shows that smaller models such as Mistral-Small and Ministral-8B achieved remarkably low inference times (0.05–0.20s), while still performing competitively in zero-shot classification. Notably, some small models approached or even matched the performance of significantly larger counterparts, indicating that latency and model size do not always correlate with performance. Ultimately, model selection should come down to how much latency, accuracy, and output robustness matter for the specific downstream classification task.

As detailed in the Results section, a paired-bootstrap analysis showed that Mistral-Small and Nemotron-70B are statistically indistinguishable on Δ-Balanced Accuracy, whereas Qwen2.5-72B lags behind. We therefore treat the former two as a single top-performance band and focus selection on latency and cost. While both Nemotron and Mistral deliver strong accuracy, they differ substantially in model scale: Nemotron is a full 70B-parameter model, while Mistral-Small is just 24B. Despite this, their inference times differ by only a few seconds. Which model is preferable may depend on the downstream task’s tolerance for model size and associated resource constraints.

## 5 Limitations

Here, we acknowledge a few limitations in our setup. Firstly, our ground truth relies on human-curated labels to determine whether a given abstract establishes a relationship between a pathogen and a disease. While necessary, this may introduce a potential bias, especially given that we are working with existing abstracts not originally written for this specific task. Errors in labeling or inconsistencies in how disagreements were resolved during the original curation process could influence both model evaluation and the conclusions we draw. Another fundamental constraint is that all models operated on abstracts and titles alone. In real-world biomedical research, important contexts that could confirm or reject a relationship between a pathogen and a disease may be buried deeper in the full text, perhaps within methods, results, or discussion sections.

By focusing solely on abstracts, we risk missing or misclassifying nuanced relationships, especially in ambiguous cases.

Secondly, we did not benchmark the LLMs against existing classical machine-learning baselines. Without those, it remains unclear whether a balanced accuracy of 0.81 represents a genuine advance over existing text classification workflows. Incorporating NLP baselines is therefore a priority for future studies.

Lastly, all models were evaluated using fixed generation parameters, with greedy decoding (temperature = 0) to ensure deterministic outputs for structured parsing. While this improved consistency, it may have limited the models’ ability to explore alternative completions. Future work could examine how different decoding settings impact performance for this task.

Looking ahead, future work could explore fine-tuning or ensemble methods to improve classification consistency. In this study, we avoided fine-tuning due to the limited dataset size and the risk of overfitting. However, with access to a larger and more diverse dataset, fine-tuning could become a viable strategy to improve model performance for our specific classification task.

## 6 Conclusion

In summary, our evaluation revealed early signs of promise in using LLMs to identify pathogen–disease relationships within PAIS literature, particularly under zero-shot settings with well-structured abstracts. While overall performance remains inconsistent, models like Mistral-Small-Instruct and Llama-3.1-Nemotron-Instruct demonstrated a strong trade-off between accuracy, inference speed, and reliability, highlighting the potential of smaller, efficient architectures in specialized classification tasks.

